# Spatially-enhanced clusterwise inference for testing and localizing intermodal correspondence

**DOI:** 10.1101/2022.04.25.489462

**Authors:** Sarah M. Weinstein, Simon N. Vandekar, Erica B. Baller, Danni Tu, Azeez Adebimpe, Tinashe M. Tapera, Ruben C. Gur, Raquel E. Gur, John A. Detre, Armin Raznahan, Aaron F. Alexander-Bloch, Theodore D. Satterthwaite, Russell T. Shinohara, Jun Young Park

## Abstract

With the increasing availability of neuroimaging data from multiple modalities—each providing a different lens through which to study brain structure or function—new techniques for comparing, integrating, and interpreting information within and across modalities have emerged. Recent developments include hypothesis tests of associations between neuroimaging modalities, which can be used to determine the statistical significance of intermodal associations either throughout the entire brain or within anatomical subregions or functional networks. While these methods provide a crucial foundation for inference on intermodal relationships, they cannot be used to answer questions about where in the brain these associations are most pronounced. In this paper, we introduce a new method, called CLEAN-R, that can be used both to test intermodal correspondence throughout the brain and also to localize this correspondence. Our method involves first adjusting for the underlying spatial autocorrelation structure within each modality before aggregating information within small clusters to construct a map of enhanced test statistics. Using structural and functional magnetic resonance imaging data from a subsample of children and adolescents from the Philadelphia Neurodevelopmental Cohort, we conduct simulations and data analyses where we illustrate the high statistical power and nominal type I error levels of our method. By constructing an interpretable map of group-level correspondence using spatially-enhanced test statistics, our method offers insights beyond those provided by earlier methods.

## 1. Introduction

A growing emphasis on extracting, synthesizing, and interpreting patterns from multiple neuroimaging modalities has simultaneously prompted innovation in methods development. One area of expanding interest is in intermodal coupling, where many researchers have sought to study and quantify correspondence between two modalities (for example, between structural and functional brain measurements). Methodological developments in this area have included statistical estimation of individual-level maps measuring intermodal coupling [1, 2, 3, 4, 5] and hypothesis tests for establishing statistical significance of intermodal associations [6, 7, 8, 9]. In this paper, we focus on the latter class of methods, examining and seeking to resolve gaps in previous methods for hypothesis testing of intermodal correspondence.

### 1.1. Intermodal correspondence testing: global testing without localization

Previous methods for hypothesis testing of intermodal correspondence aimed to address questions of whether or not correspondence exists throughout the entire brain or within pre-defined subregions of the brain. Developed by Alexander-Bloch et al. [6], the “spin test” formalized the notion of using spatially-constrained randomization for testing intermodal correspondence using group-level imaging measurements. A few years later, Burt et al. [7] proposed Brain Surrogate Maps with Autocorrelated Spatial Heterogeneity (BrainSMASH), using spatially-constrained null modeling based on reconstructing empirical variograms in spatially permuted data. More recently, we described the simple permutation-based intermodal correspondence test (SPICE), the first method to leverage participant-level as opposed to group-averaged brain maps, using a traditional permutation framework by randomly shuffling individual participants’ maps, rather than using spatially-derived null models of group-averaged brain maps [9].

Each of these recent methodological advancements for testing intermodal correspondence involves computing a single global test statistic (e.g., Pearson correlation) measuring correspondence across the entire brain (or across a pre-defined brain region, such as a functional brain network described by Yeo et al. [10])—without differentiating the degree of correspondence across the brain or within those pre-defined regions. Localizing *where* in the brain intermodal correspondence is most prominent could facilitate more nuance and interpretability in these analyses. For example, spatial localization could tell us whether structure-function correspondence across the brain is driven primarily by a subset of brain regions. While existing methods for testing intermodal correspondence are statistically powerful, they are limited in that they cannot be used to visualize or quantify local intermodal correspondence without pre-defining sub-regions where intermodal correspondence is suspected (e.g., functional networks).

### 1.2. Unique challenges in localizing intermodal correspondence

A common approach for spatially mapping and testing intermodal correspondence involves a mass-univariate analysis, where location-specific measures of association are quantified using location-specific (e.g., vertex- or voxel-wise) correlations or partial correlations. For example, in Figure 1, we map partial correlations (adjusting for age and sex) between two pairs of modalities in around 800 children and adolescents from the Philadelphia Neurodevelopmental Cohort (PNC). (A more detailed description of the PNC data and processing is provided in Section 3.1 and in Section S2 of the Supplementary Material). In such an analysis, measurements at each location in the brain are modeled separately, and *p*-values are used to “threshold out” locations in brain without significant intermodal correspondence, adjusting for multiple comparisons using the false discovery rate (FDR, as in Figure 1) or family-wise error rate (FWER). However, in smaller sample settings, mass-univariate methods suffer from low statistical power and poor replicability.

**Figure 1:**
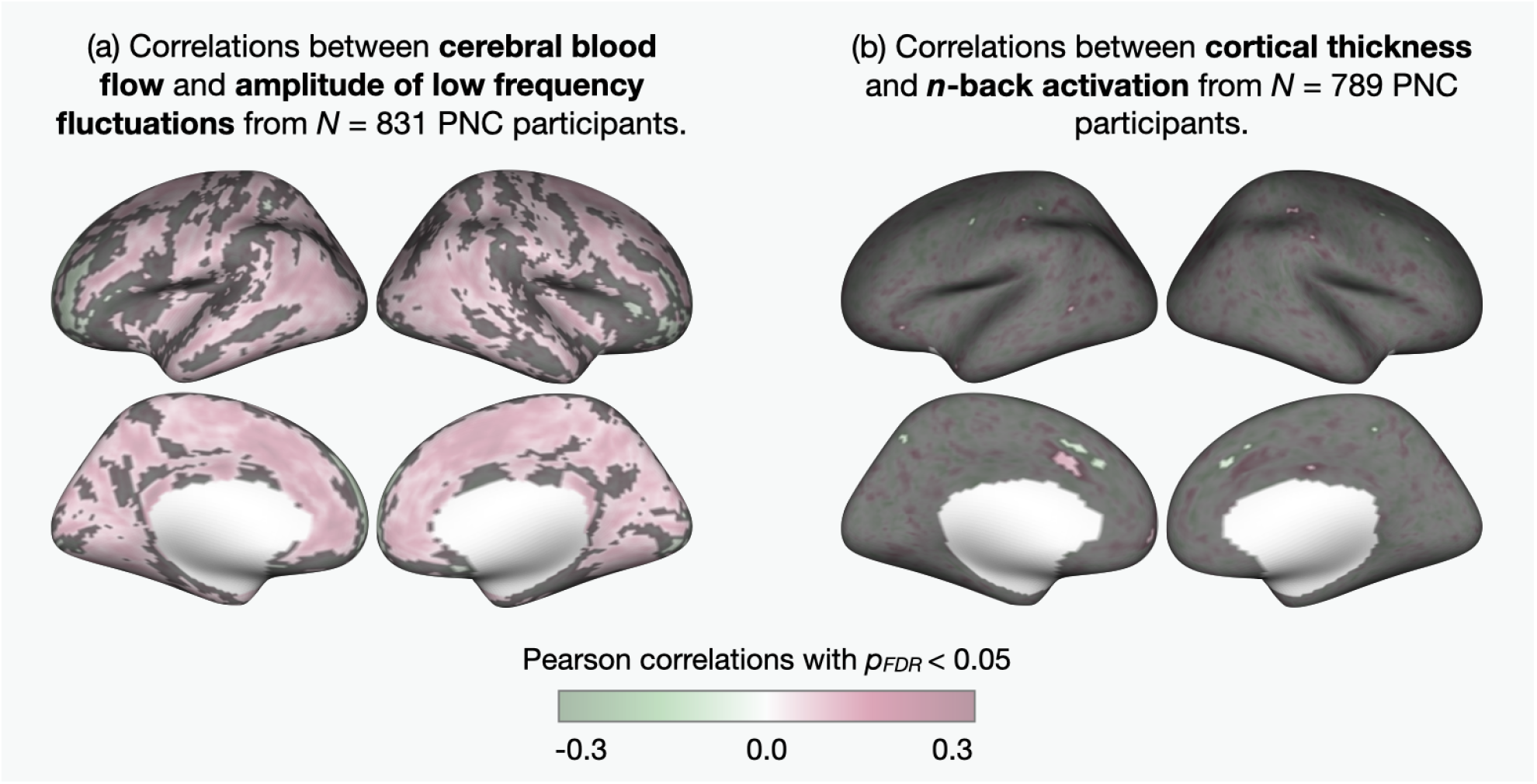
Maps of vertex-level partial Pearson correlations (adjusting for age and sex) between (a) CBF and ALFF in *N* =831 PNC participants and (b) cortical thickness and *n*-back task activation in *N* =789 PNC participants. Grayed-out areas are not statistically significant (i.e. FDR-adjusted *p*-values are greater than 0.05).

In the context of analyzing and localizing associations between a single imaging modality and a non-imaging phenotype of interest, another common approach is clusterwise inference. This is typically considered more powerful than mass-univariate, since univariate test statistics are aggregated in the spatial domain [11, 12, 13]. Inference using clusterwise inference approaches uses permutation across subjects, controlling FWER at the nominal level [14].

A fundamental limitation inherent to using univariate test statistics (including mass-univariate or standard clusterwise inference methods) is the absence of modeling complex spatial autocorrelation patterns inherent in neuroimaging data. As an illustration, we consider Figure 2, where we quantify and visualize the underlying spatial autocorrelation structures in four different imaging modalities from the PNC by using empirical variograms. Here, it is clear that there is heterogeneity in brain-wise spatial autocorrelation exhibited across different imaging modalities. We therefore anticipate that between-modality variability will be a critical issue in localizing intermodal correspondence. Importantly, such heterogeneity in spatial autocorrelation in multimodal data may interfere with scientific interpretation or lead to a severe loss of power in the context of applying univariate statistical methods.

**Figure 2:**
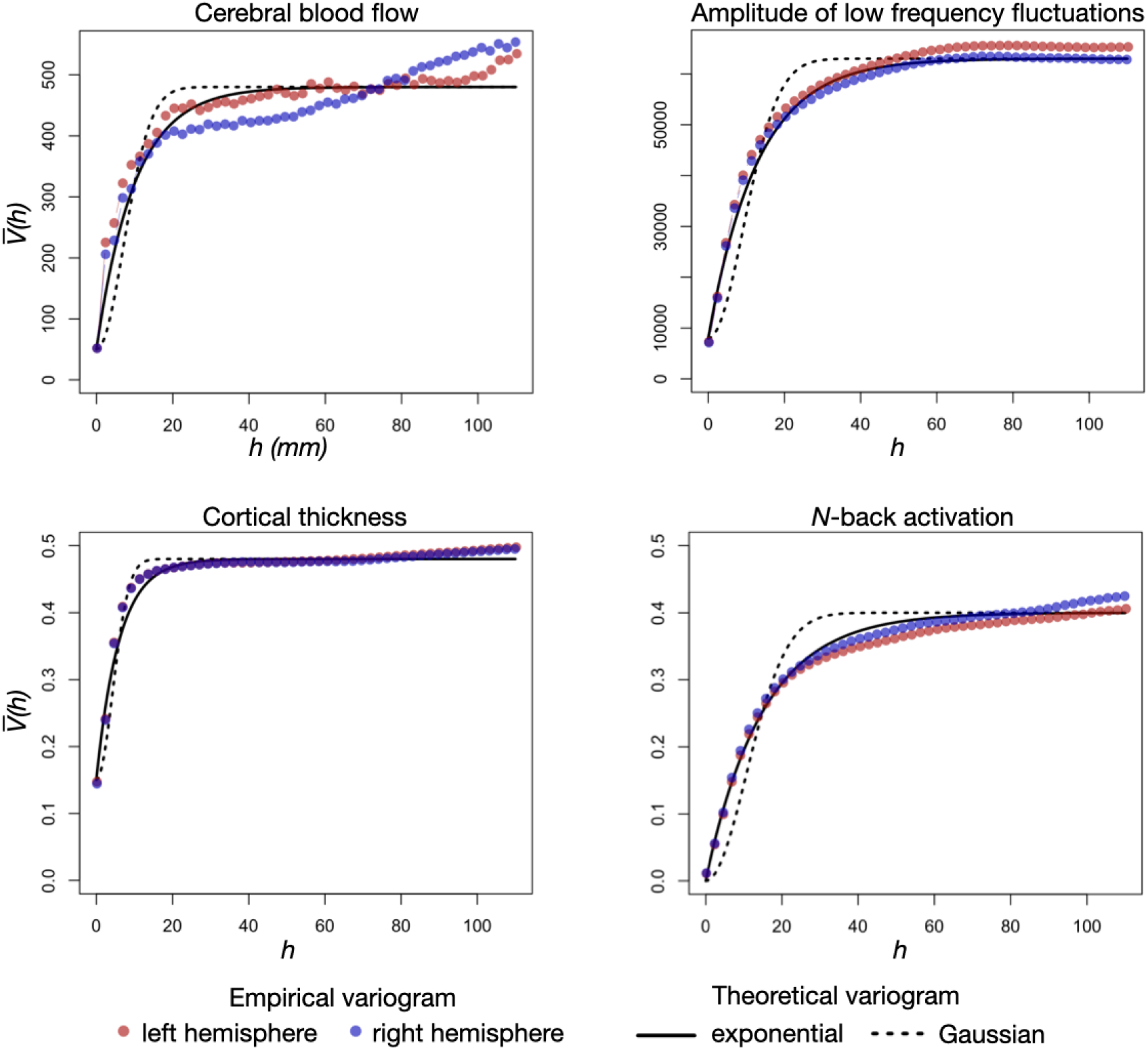
Group-averaged empirical variograms in four neuroimaging modalities from the Philadelphia Neurodevelopmental Cohort (PNC) [15], described in Section 3 and Section S2 of the Supplementary Material. The empirical variogram provides a measure of how strong the spatial autocorrelation between different points in an image depends on the distance *h* between those points [16]. For each imaging data modality from the PNC, we compute the empirical variogram for each participant (*V*_*i*_(*h*) for the ith participant across different *h*) and then average it across subjects to obtain group-averaged empirical variogram 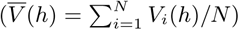. We use geodesic distance to measure distances (in millimeters) between vertex pairs on the cortical surface. For comparison, we also plot hypothetical theoretical variograms with exponential (solid line) and Gaussian (dotted line) structures, which informs the parametric assumptions used in our models for spatial adjustment, discussed in Section 2.2.

### 1.3. Motivating clusterwise inference with modality-specific modeling of spatial autocorrelation to test and localize intermodal correspondence

Recognizing the unique challenge of leveraging spatial information in an analysis of multiple imaging modalities with inherently different underlying spatial structures, we propose CLEAN-R, a novel method that combines insights from Weinstein et al. [9] and Park and Fiecas [13] to address this important gap in existing methodology. The general framework for our proposed method is as follows (described in greater detail in Section 2). First, we remove nuisance covariate effects and spatial variations separately for each modality, using an approach following Park and Fiecas [13] in the context of the general linear model (GLM) for a single imaging modality. Next, we aggregate location-specific test statistics within optimized cluster-level radii to construct enhanced statistics that spatially map intermodal correspondence throughout the brain. In addition, we use the permutation framework of Weinstein et al. [9], controlling the type I error rate and FWER at the nominal level. Finally, we propose a statistical framework to localize brain regions where there are significant group differences in intermodal correspondence. Our method is inherently computationally efficient and can be implemented using our publicly available software (www.github.com/junjypark/CLEAN).

In Section 3, we apply CLEAN-R to the data from the large-scale PNC study to demonstrate our method in practice and compare its performance with other approaches to hypothesis testing and spatial localization of intermodal correspondence. Using data from the PNC, we construct data-driven simulations under both null and non-null cases, and show that CLEAN-R controls FWER and exhibits high statistical power for both global hypothesis testing and spatial localization of intermodal correspondence. Importantly, we find that clusterwise inference without modeling modality-specific autocorrelation suffers from poor replicability across different sub-samples from the PNC, which highlights the added value of spatial adjustment used in CLEAN-R.

## 2. Methods

### 2.1. Notation and null hypothesis

We consider a dataset consisting of participant-level measurements from two different neuroimaging modalities. For example, **x**_*i*_ could be an image of brain structure for the *i*th participant, and **y**_*i*_ a measure of brain function for the same participant. We assume that data from both modalities are available for all *N* study participants, and that each modality consists of *V* measurements (e.g., vertices on the cortical surface). Letting *x*_*i*_(*v*) and *y*_*i*_(*v*) be the *i*th individual’s measurements from both modalities at location (or vertex) *v*, we define **x**(*v*) = (*x*_1_(*v*), …, *x*_*N*_ (*v*))^*′*^ and **y**(*v*) = (*y*_1_(*v*), …, *y*_*N*_ (*v*))^*′*^ to be vectors of data derived from each modality at a single image location *v* across all *N* study participants, where *v* = 1, …, *V*. We can also represent participant-level images (across all locations) as vectors **x**_*i*_ = (*x*_*i*_(1), …, *x*_*i*_(*V*))^*′*^ and **y**_*i*_ = (*y*_*i*_(1), …, *y*_*i*_(*V*))^*′*^.

We express our global null hypothesis that there is no correspondence between the two modalities as follows:

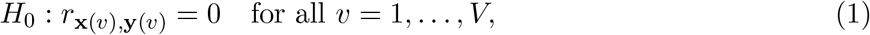

where *r*_**u**,**v**_ is the Pearson correlation between vectors **u** and **v**. In the presence of participant-level nuisance covariates (e.g. age and sex), we express the global null hypothesis in terms of partial correlations, by replacing **x**(*v*) and **y**(*v*) with **x**^⋆^(*v*) and **y**^⋆^(*v*), respectively, where **x**^⋆^(*v*) and **y**^⋆^(*v*) are the estimated residuals we obtain after regressing out nuisance covariates from **x**(*v*) and **y**(*v*) (separately for each *v*), while preserving the same dimension as the original data. If *H*_0_ is true, our interpretation is that there is no location *v* in the entire brain where there is an association between **x**(*v*) and **y**(*v*).

### 2.2. CLEAN-R: <u>C</u>lusterwise inference <u>le</u>veraging spatial <u>a</u>utocorrelation in <u>n</u>euroimaging for testing intermodal correlations <u>r</u>

#### 2.2.1. Leveraging modality-specific spatial autocorrelation

As illustrated in Figure 2, there is substantial variability in the underlying spatial structures of different neuroimaging modalities, and we anticipate this will have implications for how to best handle spatial autocorrelation in the context of intermodal association studies. Therefore, we propose accounting for differences in the spatial autocorrelation structure of each modality before conducting intermodal correspondence testing. To do this, we extend the framework in modeling brain-wise spatial autocorrelation from Park and Fiecas [13] to account for underlying differences between the spatial structures across modalities.

Park and Fiecas [13] first explicitly model the spatial autocorrelation structure of participant-level images by using a spatial Gaussian process. Analogously, we make parametric assumptions about the covariance structure of 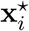 and 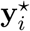 to facilitate adjustment for spatial autocorrelation. Specifically, we assume 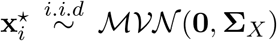 and 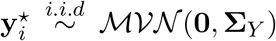, where the covariances decompose into the sum of their respective spatial and non-spatial variances as follows:

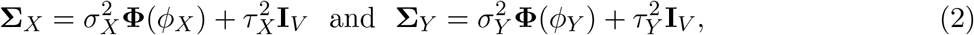

where **Φ**(*ϕ*_*X*_) and **Φ**(*ϕ*_*Y*_) are the assumed parametric spatial autocorrelation functions and 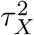 and 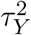 are the nugget effects measuring residual non-spatial variability in **x**_*i*_ and **y**_*i*_. After determining appropriate parametric assumptions, we estimate parameters in Equation (2) using the method-of-moments approach described by Park and Fiecas [13].

It is important to emphasize that the use of parametric assumptions in our proposed method must be justified through exploratory analyses using empirical variograms, as we have illustrated in Figure 2, and that using an incorrect spatial autocorrelation function (SACF) might result in loss of power. In particular, in Figure 2, we directly compare group-averaged empirical variograms from four different neuroimaging modalities (blue and red dots) to hypothetical theoretical variograms based on exponential (solid lines) and Gaussian or squared exponential (dotted lines) parameterizations. We observe a close alignment between the theoretical exponential variograms and empirical variograms for all four modalities; therefore, we consider an exponential covariance structure to be a reasonable parametric assumption—that is, we assume the (*v, v*^*^)th element of the spatial auto-correlation functions, **Φ**(*ϕ*_*X*_) and **Φ**(*ϕ*_*Y*_), are exp(−*ϕ*_*X*_ × *d*_*v,v\**_) and exp(−*ϕ*_*Y*_ × *d*_*v,v\**_), respectively, where *d*_*v,v\**_ is the geodesic distance between locations *v* and *v*^*^. Emphasizing that the current parametric assumptions are based specifically on Figure 2, we advise future researchers to conduct similar exploratory visualizations before making any parametric assumptions and proceeding with subsequent steps.

It is also worth noting that Equation (2) provides an explicit random effects formulation of the Gaussian process, which is equivalent to setting 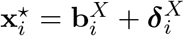 where 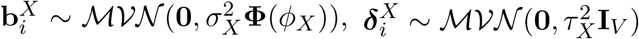 and 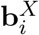 and 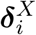 are independent (and similarly, setting 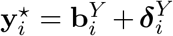 where 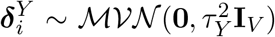 and 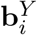 and 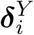 are independent). As 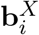 and 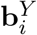 represent the portion of 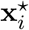 and 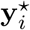, respectively, explained solely by spatial autocorrelation and not by any other factors that are truly of interest (e.g., intermodal correspondence) we consider these to be unwanted or “nuisance” sources of variation and replace them with 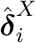 and 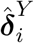, defined for the *i*th subject by

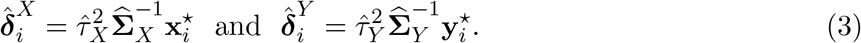

Thus, we characterize 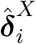 and 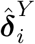 to be linear transformations of 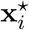 and 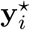, respectively, which have been adjusted for their unique underlying spatial autocorrelation structures.

We note that when the spatial parameters above are estimated via restricted maximum likelihood (REML), 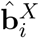 and 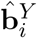 have the desirable property of being the empirical best linear unbiased predictions (eBLUP) of the spatial variations, making 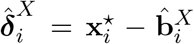 and 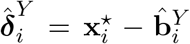 good predictions of the non-spatial variation from 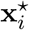 and 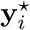, respectively [17]. In practice, the method-of-moments approach we use for estimation yields estimates that are very close to those from REML, but are much more computationally efficient.

Lastly, while inverting 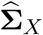 and 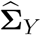 (Equation (3)) can be computationally intensive, especially for high-dimensional imaging data, we approximate the inverse of 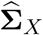 and 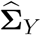 with 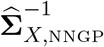 and 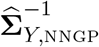, respectively, which we obtain via a nearest-neighbor Gaussian process (NNGP) [18, 19]. NNGP is a computationally efficient way to obtain a close approximation of the inverse of the parametric spatial covariance matrix using information from a subset of neighbors, which was also adopted in CLEAN [13].

#### 2.2.2. Proposed spatially-enhanced clusterwise inference for testing intermodal correspondence

In this section, we describe implementing clusterwise inference after adjusting for modality-specific spatial variations to improve the sensitivity of localized signals. In contrast to existing clusterwise inference methods commonly used in neuroimaging, the proposed method utilizes spatial information in two ways: (i) modeling and adjusting for noise (Section 2.2.1) and (ii) aggregating test statistics from nearby locations (current section). Briefly, (ii) involves computing vertex-level statistics (using 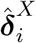 and 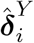 from (i)), and then combining these to obtain cluster-level statistics. The steps for computing spatially-enhanced vertex-level test statistics are summarized in the remainder of this section and in pseudocode included in Section S1 of the Supplementary Material (Algorithm S1.1).

First, we compute the vertex-level test statistics using the Fisher-transformed Pearson correlation between the two modalities at each location:

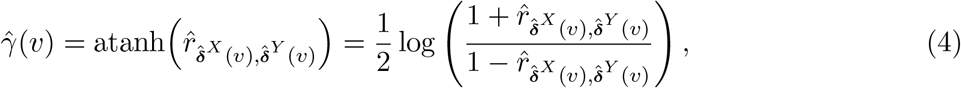

where 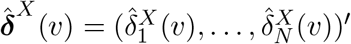 and 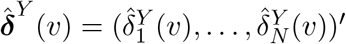 (from Equation (3)). We note that 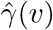 is approximately normally distributed with mean 0 under the null hypothesis.

Second, we compute cluster-level test statistics using 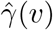 in Equation (4) as a building block. Let *N*_*h*_(*v*) denote the set of vertices that fall within radius *h* from a central vertex *v. N*_*h*_(*v*) is initially defined for each location and different possible radii. Figure 3 illustrates a single vertex on the cortical surface (black dot) and its neighbor set *N*_*h*_(*v*) with different values of *h*. Suppose for each vertex, we consider clusters defined by *m* different radii, *h*_1_, …, *h*_*m*_. Assuming *h*_1_ = 0, 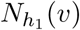 is the cluster consisting of only the central vertex *v*. Thus, the correspondence measure for this cluster is equal to 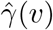, and the test statistic for location *v* for a cluster with radius up to *h*_*j*_ (*j* = 2, …, *m*) are computed as the standardized sums of 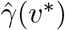 for all 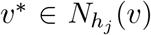. It motivates the construction of a unified cluster-level test statistic for *v* by

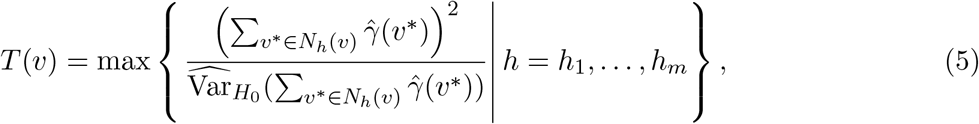

so that the optimal cluster size for a given vertex can be determined by the magnitude of cluster-level test statistics with different radii. Accordingly, a global statistic for testing *H*_0_ (Equation (1)) is

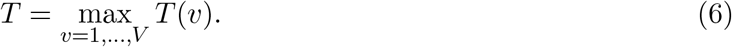

Following Park and Fiecas [13], in the analyses below we consider *h*_1_ = 0 to *h*_*m*_ = 20mm in increments of 1mm (as in Figure 3) as a default; however, we explore other maximum radii *h*_*m*_ in the Supplementary Material (Section S3).

**Figure 3:**
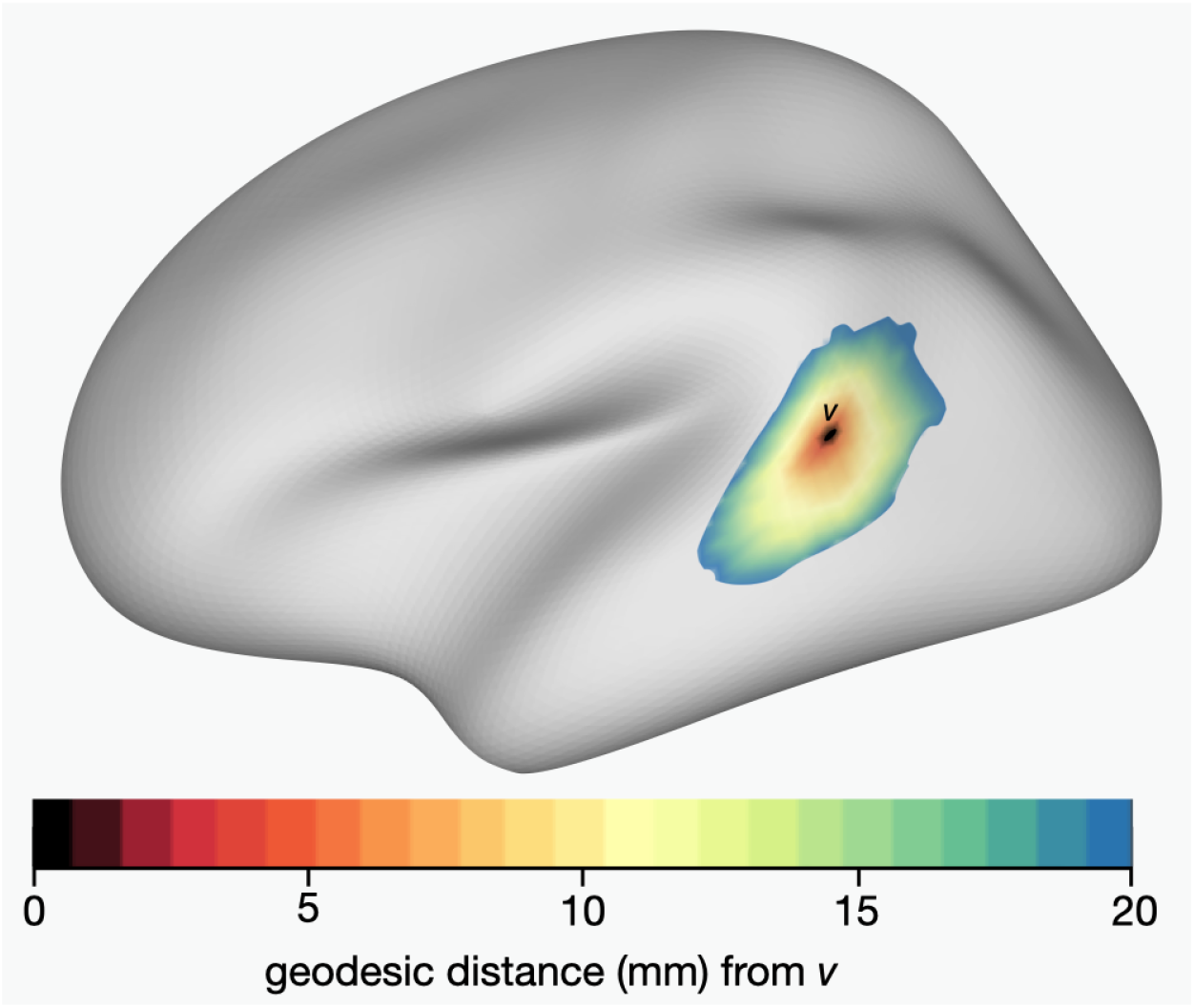
Example clusters surrounding vertex *v* (black dot) in the left hemisphere, with a maximum radius of 20mm on the cortical surface (measured as a geodesic distance). Our proposed method involves defining candidate clusters for each vertex, with different maximum radii ranging from 0 to 20mm. As we describe in Section 2.2, we compare test statistics that have been aggregated across image locations within different sized clusters surrounding each vertex to determine the optimal sized cluster to enhance the test statistic measuring intermodal correspondence at each location.

An important step in conducting statistical inference with the proposed statistic is the selection of a threshold (*t*_*α*_), or critical value, to control FWER at *α* (e.g. 0.05). Once a brain-wise threshold is set, we can localize significant intermodal correspondence at vertices whose *T* (*v*) exceeds the threshold. Specifically, we propose permuting data across participants within one modality, following Weinstein et al. [9]. Because our null hypothesis is expressed in terms of within-participant intermodal correlations, randomly shuffling participants within one modality destroys any associations between two modalities derived from the same individual; thus, an empirical null distribution made up of inter-individual correlations is appropriate for testing the null hypothesis (Equation (1)). In the presence of participant-level covariate information, we preserve the order of the covariates and the shuffled modality in each permutation because our hypothesis addresses residual (partial) correlation of two modalities after regressing out non-imaging covariates. For the *k*th permutation (*k* = 1, …, *K*), we construct *T* ^(*k*)^(*v*) using Equation (5) then construct its maximum across all vertices,

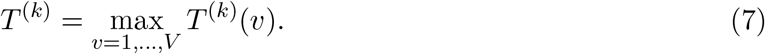

and choose the threshold *t*_*α*_ bt the (1 − *α*) × 100th percentile of the empirical null distributions constructed by *T* ^(1)^, …, *T* ^(*K*)^.

### 2.3. Testing and localizing group differences in intermodal correspondence

In this section, we extend CLEAN-R for testing and localizing group differences in intermodal correspondence, utilizing the advantages of clusterwise inference and autocorrelation modeling involved in CLEAN-R. In this setting, we restate the null hypothesis (Equation (1)) in terms of the difference in intermodal correlations between two groups (say, groups *A* and *B*) at vertex *v* (e.g., 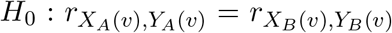 for all *v*). Let 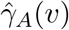 and 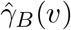 denote separately estimated location-specific association measures (as in Equation (4)) for each group. To test and localize differences in intermodal correspondence these two groups, we replace 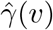 in Equation (5) with 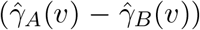. The spatially-enhanced statistic used for testing differences between groups at vertex *v* is analogously given by

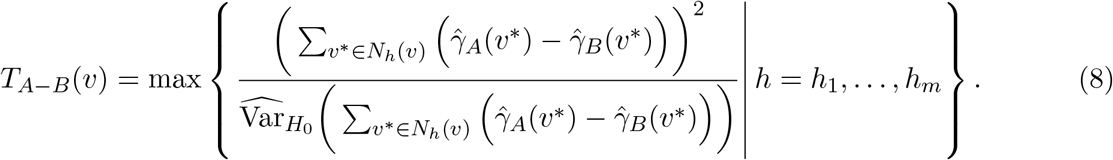

After permuting participants separately within each group, an empirical null distribution for both global and local tests of group differences in correspondence is obtained analogously to (7); that is,

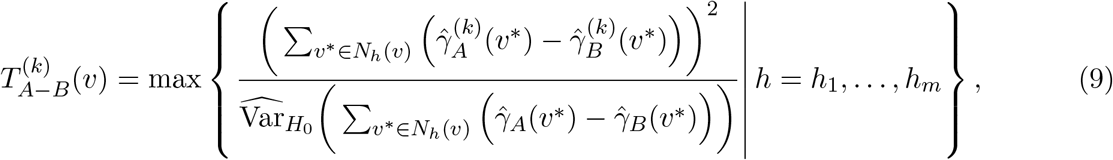

and similar to *T* ^(*k*)^ (Equation (7)), 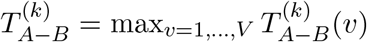.

### 2.4. Comparison to other methods for testing intermodal correspondence

In this section, we compare different approaches in testing intermodal correspondence, including BrainSMASH [7], the spin test [6], SPICE [9], and classical approaches such as mass univariate and clusterwise inference without spatial adjustment.

#### 2.4.1. Permutation null versus spatial null

Although all methods we consider use some form of permutation or spatial randomization to generate null maps, they are fundamentally different. We summarize key distinctions between permutation and spatial null models in Figure 4. The spin test and BrainSMASH take different approaches to spatial randomization to generate null maps. Both methods first use a group-averaged map in each modality. Then, the spin test repeatedly rotates an imaging modality (projected onto a spherical surface) and computes test statistics measuring correspondence between the “rotated” map and original second map to generate an empirical null distribution. BrainSMASH randomly shuffles vertex-level (or parcel-level) measurements across the brain (thus destroying both the spatial autocorrelation structure as well as any genuine intermodal associations), then applies Gaussian smoothing to generate surrogate maps with similar empirical variograms as the original map.

**Figure 4:**
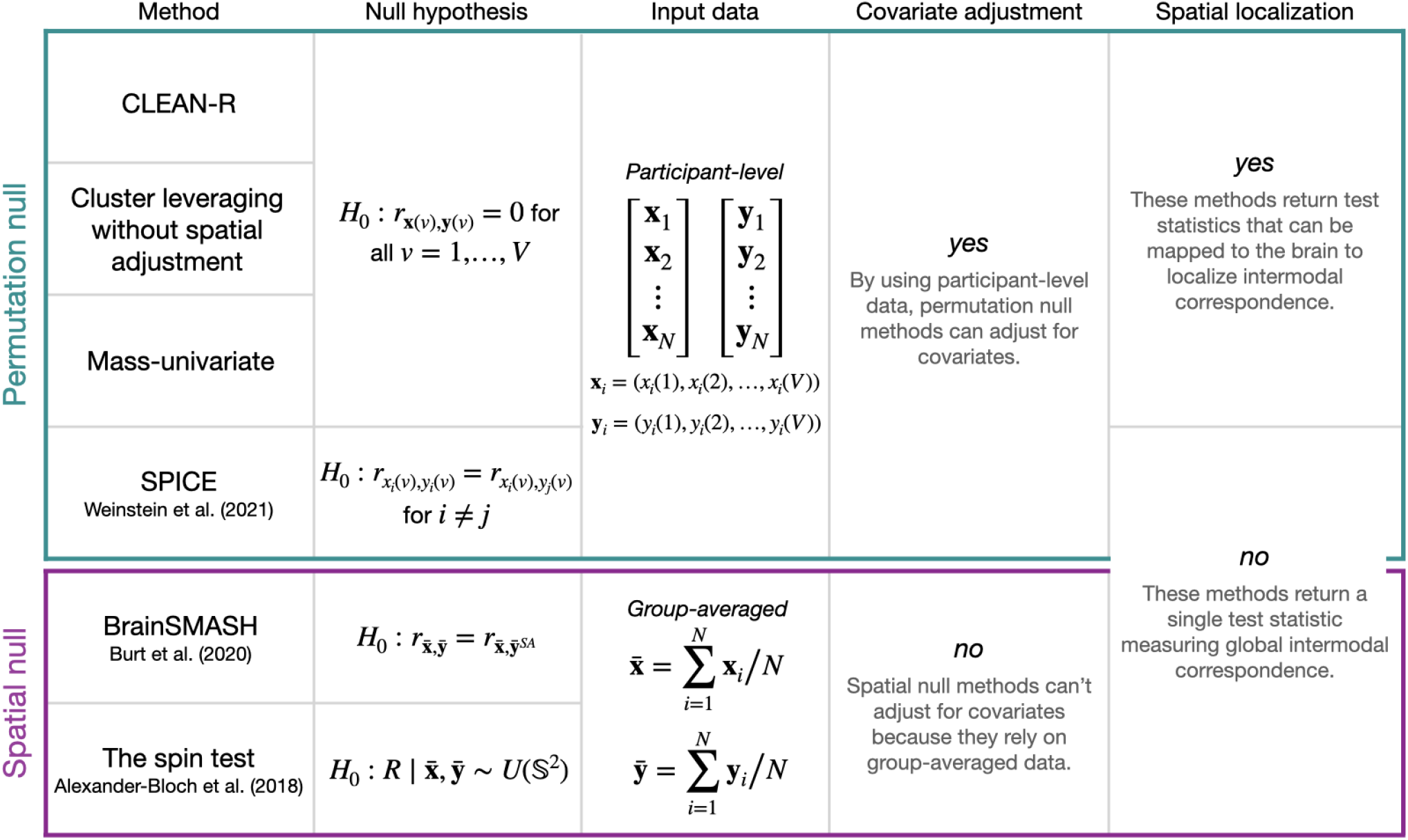
Summary and comparison of CLEAN-R to other methods for testing intermodal correspondence. We distinguish between two broad classes of methods for hypothesis testing of intermodal correspondence based on the types of null models used: either permutation null or spatial null. While methods using a permutation null approach leverage participant-level data to generate an empirical null distribution based on between-participant intermodal associations, spatial null models rely on group-level brain measurements. While a comparison across all these methods is warranted due to the popularity of the two spatial null models by Alexander-Bloch et al. [6] and Burt et al. [7], differences in the input data and capabilities (e.g., covariate adjustment and spatial localization) of these methods restrict which methods can be considered in the analyses described in Section 3. *Note*: In the null hypothesis for BrainSMASH, 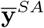 denotes a spatial autocorrelation (SA)-preserving “surrogate” brain map, which Burt et al. [7] use to evaluate whether any observed correlations between 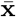 and 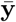 are genuine or are merely driven by spatial autocorrelation.

While BrainSMASH is an important development in hypothesis testing involving neuroimaging data, several limitations warrant further consideration, including the implicit assumption that the spatial autocorrelation structure of both modalities is similar, and that stationarity can be assumed throughout the brain, as well as within anatomical subregions and functional networks when more localized tests of intermodal correspondence are considered. We also point out that, by considering a group-averaged map in each modality, neither the spin test nor BrainSMASH can adjust for confounders or other nuisance covariates, such as age or sex. For example, if a group-averaged map is constructed after regressing out confounding variables, then it just becomes zero for all locations (due to properties of GLM residuals), making an evaluation of correspondence infeasible.

In contrast, permutation null methods that shuffle images from a single modality across participants (including CLEAN-R, SPICE, and mass-univariate methods) provide a clear null hypothesis defined non-parametrically. By using participant-level rather than group-averaged data, we can easily adjust for confounders through residualization. Also, these methods can be implemented in an efficient and scalable manner. Based on our implementation, BrainSMASH and other methods involving spatial null models summarized by Markello and Misic [8], are much more computationally intensive.

Because of differences in null hypotheses and underlying assumptions, we note that direct comparisons with the spatial null methods (the spin test and BrainSMASH) may not always be possible. Therefore, in our analysis in Section 3, we primarily focus on comparing methods based on the permutation null mapping, and subsequently discuss results from the spatial null methods in Section 3.2.3.

#### 2.4.2. On true spatial autocorrelation

A key distinction between CLEAN-R and existing methods for clusterwise inference is its spatial autocorrelation modeling. Despite their wide utility, Figure 2 suggests that in the presence of spatial autocorrelation, existing clusterwise inference methods are misspecified because these are based on univariate statistics, which ignore spatial autocorrelation. We note that a parametric assumption of brain-wise stationarity in CLEAN-R may also be misspecified, because empirical variogram implicitly assumes spatial stationarity. Still, we believe that the proposed approach is closer in approximating the true underlying autocorrelation structure, which can be validated through evaluation of statistical power.

### 2.5. Software and computational efficiency

CLEAN-R can be implemented using software available at www.github.com/junjypark/CLEAN. A detailed algorithm is also summarized in Section S1 of the Supplementary Material (Algorithm S1.1). Importantly, the proposed permutation strategy is inherently computationally efficient because permuting information across participants (along with corresponding covariate information) does not affect the variance component estimates of the original model. Therefore, permutation is not limited to shuffling 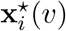 but is also applicable to 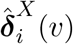 from the proposed model.

For example, applying CLEAN-R to a sample of *N* =50—with nearly 10,000 vertices in each hemisphere for each person and using 10, 000 permutations—takes under 14 minutes. This includes the time to fit four covariance regression models (one for each modality in each hemisphere), and it can be even more efficient with parallelization, which is also supported by our software.

## 3. Simulations and data analysis

### 3.1. Multimodal neuroimaging data from the Philadelphia Neurodevelopmental Cohort

The Philadelphia Neurodevelopmental Cohort (PNC) is a large-scale research initiative led by the Brain Behavior Laboratory at the University of Pennsylvania and Children’s Hospital of Philadelphia, which recruited *N* =9,498 children and adolescents ages 8-21 years old who consented or received parental consent to participate [20]. A subset of 1,601 participants from the PNC underwent structural and functional MRI scans at the Hospital of University of Pennsylvania in the same Siemens TIM Trio 3T scanner according to protocols described in detail by Satterthwaite et al. [15]. In this paper, we consider data from four different neuroimaging modalities to examine intermodal associations involving two different pairs of modalities in the PNC: (i) cerebral blood flow (CBF) and amplitude of low frequency fluctuations (ALFF) and (ii) cortical thickness and *n*-back activation, discussed below in Section 3.1.1.

Participant-level images from each modality were collected and processed according to standardized protocols described in Satterthwaite et al. [15]. A brief summary of data collection, processing, and quality control is also included in Section S2 of the Supplementary Material. To represent participant-level images on the cortical surface, we use Freesurfer version 5.3 to resample voxelwise measurements from each imaging modality to the fsaverage5 atlas [21]. Resulting images consist of 10,242 vertices per hemisphere per participant. For all analyses, the number of vertices in the left and right hemispheres reduces to *V* =9,354 and *V* =9,361, respectively after we exclude portions of each hemisphere corresponding to the medial wall, a byproduct of working with a surface-based representation of brain images [22]. Vertices in the medial wall are then added back for visualization purposes, where we used the `fsbrain` package in `R` [23].

In each analysis, we exclude participants who did not have data available, did not meet quality control criteria for both imaging modalities, or had other health-related reasons for exclusion (e.g., taking psychoactive medications or previous psychiatric hospitalization). For CBF/ALFF, our cohort consists of *N* =831 individuals (mean age 15.62 (SD 3.36), 478 (57.52%) females), while for cortical thickness/*n*-back, we analyzed data from *N* =789 individuals (mean age 14.96 (SD 3.19), 446 (56.53%) females). In permutation null methods for testing intermodal correspondence, we first fit a linear regression model at each image location (i.e., at 9,354 locations in the left hemisphere and 9,361 in the right hemisphere), where we include age and sex as covariates and subtract out their estimated effects in order to adjust for these (non-spatial) sources of variation. (Note: in Section 3.3 where we evaluate sex differences in correspondence between CBF and ALFF, we adjust for age but not sex).

#### 3.1.1. *H*_0_ *in context*

Increased blood flow to different areas of the brain allows for neuronal activity in those regions, and relating these two processes through the comparison of neuroimaging modalities can help us to better understand this innate biological process, how it changes throughout development, and how it differs across people. Using arterial spin labeling (ASL) to quantify cerebral blood flow (CBF) throughout the brain and the amplitude of low-frequency fluctuations (ALFF) in resting-state blood-oxygen-level-dependent (BOLD) fMRI as a proxy for neuronal activity, in this study we examine the relationship between CBF and ALFF among PNC participants. Recently, Baller et al. [24] documented the evolution of individual-level “coupling” between CBF and ALFF over the course of childhood and adolescence among PNC participants. These authors found significant coupling between CBF and ALFF at the participant-level, both throughout the cortex and in some specific regions, noting differences in the degree of coupling throughout the cortex among different age groups and by sex. In the current work, we leverage participant-level data to map, test, and localize intermodal correspondence between these two fMRI-derived brain maps at the group-level (in contrast, Baller et al. mapped these relationships for individual participants). Adding to Baller et al.’s findings, we expect to reject the null hypothesis in support of the notion that blood flow and neuronal activity are strongly related. We will further be able to localize what subregions of the brain are most strongly implicated in the CBF/ALFF relationship.

It is also informative to consider a setting in which we do not expect to see strong intermodal associations. Thus, we also test correspondence between cortical thickness and *n*-back task activation. We also compared these modalities in our previous work [9]; however, in our current work, we now leverage spatial information with CLEAN-R and may detect signal that is highly localized to specific areas of the brain, which was not possible in our earlier work.

In Section 1, we considered vertex-level Pearson correlations from the full samples of both imaging modality pairs from an exploratory analysis using the PNC data (Figure 1). As these correlations were estimated in relatively large samples of around 800 participants each, it is reasonable to consider Figure 1 as somewhat of a “ground truth,” providing insight into which regions we might hope to find evidence for localized intermodal correspondence in analyses in smaller samples. It also allow us compare different methods in terms of replicability in smaller samples.

### 3.2. Data-driven simulation study in the Philadelphia Neurodevelopmental Cohort study

We conduct a set of data-driven simulation studies using real data from the PNC to evaluate and compare the power and false positive rate of CLEAN-R in to other permutation-based methods for testing intermodal correspondence. This type of non-parametric resampling-based approach to simulation is often used in settings where the target data of interest are inherently complex and would be difficult if not impossible to emulate through parametric simulations. Resampling-based simulations have been used in previous studies validating methods in neuroimaging [25, 26], as well as numerous other applications in biomedical research (e.g., pharmacoepidemiology [27] and RNA-seq data analysis [28]).

In our simulations, we consider two different sample sizes (*N* = 50 and *N* = 100). In each design, we sample *N* participants randomly (without replacement) based on the null and alternative hypotheses, and repeated this 1000 times, resulted in 1000 simulated datasets in total. To evaluate the false positive rate of each method, in each sample, we ensure that the subject’s IDs in one modality do not match with the IDs in the other modality. To evaluate power, each pair of modalities is derived from the same individual (i.e., we do not mis-match IDs in each pair of modalities).

In summary, our entire simulation study considers eight scenarios (with 1000 simulations per scenario), which are characterized by a combination of the following:

1. The pair of modalities under investigation: (a) CBF and ALFF (widespread signal) or (b) cortical thickness and *n*-back activation (sparse signal).
2. The number of participants to be sampled (without replacement) from the full group of PNC participants (after exclusion criteria): (a) *N* = 50 or (b) *N* = 100
3. Hypotheses: (a) null (for type I error rate) and (b) alternative (for power).

Due to the design of our simulation studies, we focus on comparing the methods that use a permutation null to evaluate type I error rates and power (SPICE, mass univariate, and CLEAN-R with and without spatial autocorrelation modeling). Results from these simulations are presented below in Section 3.2.1). When we evaluate localization in Section 3.2.2, we do not consider SPICE because it does not provide location-specific measures of intermodal correspondence (rather, it can only be used for global testing). Lastly, we note that the proposed simulation method does not translate well for evaluating the two spatial null modeling (the spin test [6] and BrainSMASH [7]). However, where appropriate, we discuss comparisons with these methods in Section 3.2.3.

#### 3.2.1. Simulation results: type I error and power

Simulation results for our global tests of intermodal correspondence are included in Figure 5. All four methods show nominal 5% type I error rates for testing CBF/ALFF and cortical thickness/*n*-back correspondence in simulation settings where the null hypothesis is true. Statistical power for tests of correspondence between CBF and ALFF suggest that CLEAN-R performs similarly to the SPICE test and outperforms the spatially un-adjusted clusterwise approach and the mass-univariate approach. Because the SPICE test does not provide localization of intermodal coupling, CLEAN-R is especially appealing as it achieves high statistical power and also has the added benefit of spatial localization, which we illustrate and discuss below.

**Figure 5:**
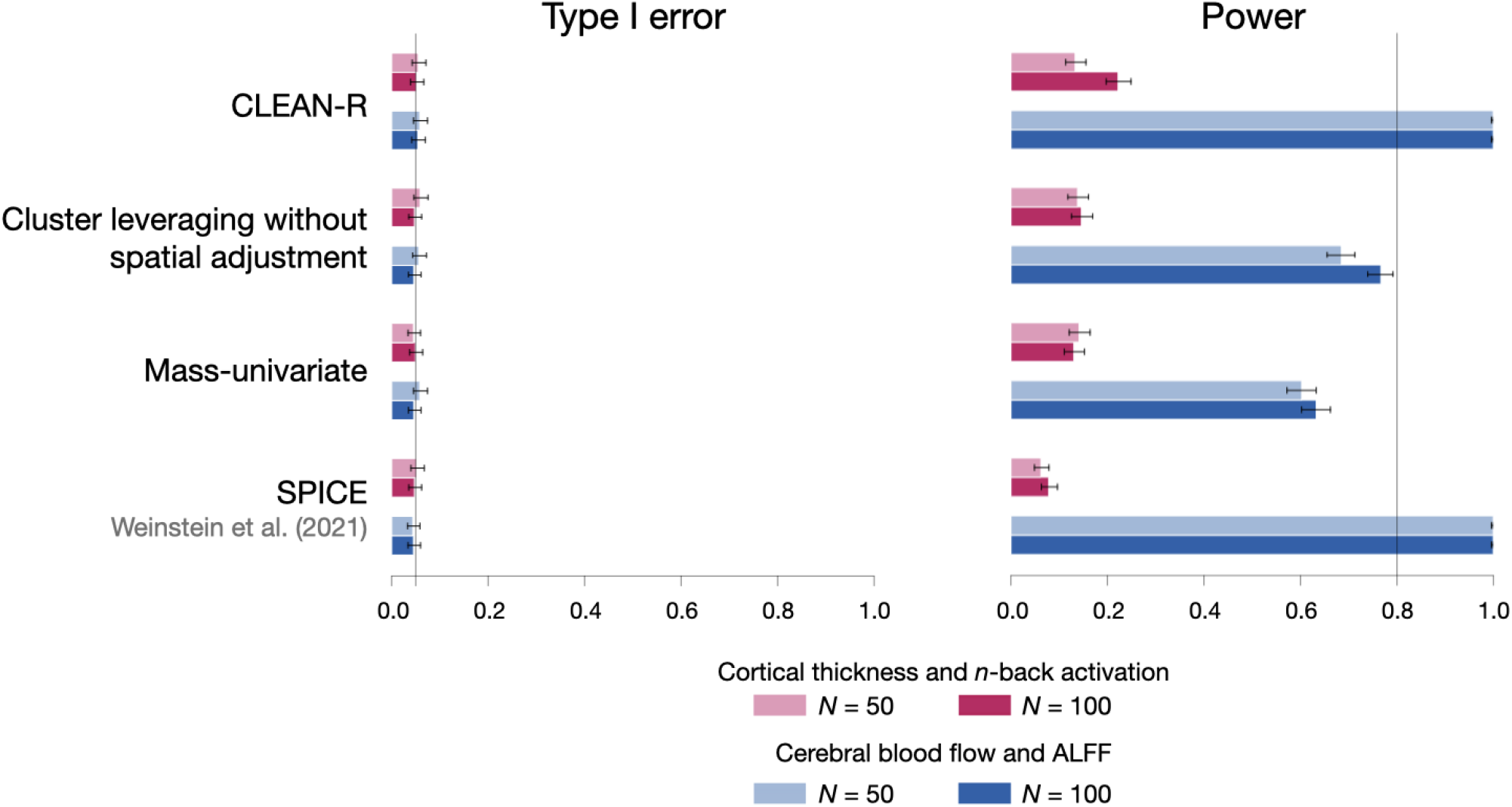
Rates of rejecting global null hypotheses from four permutation-based approaches for testing intermodal correspondence in data-driven simulation studies using data from the Philadelphia Neurodevelopmental Cohort (PNC). From 1,000 simulations conducted under the null hypothesis (using 2,000 permutations per test) for each pair of modalities and each sample size, type I error rates are the proportion of simulations where *H*_0_ was rejected. From 1,000 simulations conducted using within-participant data for each pair of modalities and sample size, power (or the rate of rejecting *H*_0_ in the observed data) is the proportion of simulations where *H*_0_ was rejected. 95% binomial confidence intervals (black segments) are provided, showing that the type I error rates in every setting are close to the nominal level of 0.05.

For tests of cortical thickness and *n*-back, no method exhibits high power. However, power for CLEAN-R was higher than any other method. Given the small regions exhibiting signal in the full sample correlation maps (Figure 1b), correspondence between cortical thickness and *n*-back may be localized to extremely small regions, making statistical power lower for methods that do not leverage enhanced cluster-level statistics as in CLEAN-R.

#### 3.2.2. Simulation results: localization of intermodal correspondence

Figure 6 maps statistical power from each of the three methods that involve estimating vertex-level test statistics (i.e. *T* (*v*) from Equation (5)). For tests of correspondence between CBF and ALFF, CLEAN-R is the only method to outline clear regions of strong statistical power for tests of correspondence, adding to our understanding of anatomical regions and functional networks that may be driving significant results in global correspondence tests. For tests of correspondence between cortical thickness and *n*-back, no method delineates specific areas of the brain with high statistical power.

**Figure 6:**
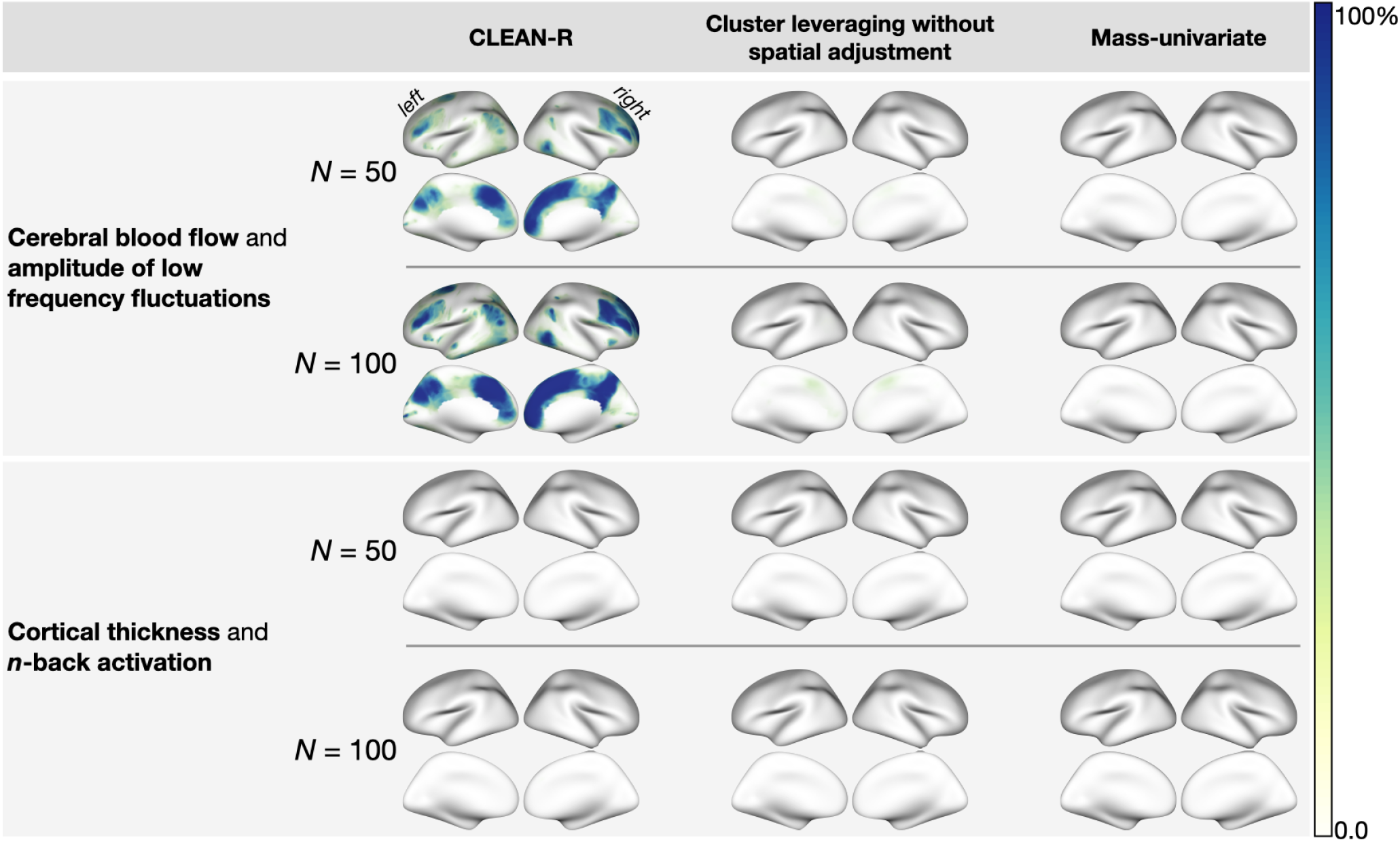
Statistical power from three different methods mapped throughout cortex. Map values between 0 and 100% indicate the proportion of simulations where *H*_0_ was rejected at a particular location. Power for global tests of coupling between each pair of modalities according to each method are provided in Figure 5. At each vertex *v*, statistical power is the proportion of the 1000 simulations where the vertex-level test statistic (e.g. *T* (*v*) as in Equation (5)) exceeds the FWER-controlling threshold *t*_*α*_.

In the Supplementary Material, we provide several additional comparisons to the results presented in Figure 6. In Figure S3.1, we evaluate the impact of the choice of the maximum radius *h*_*m*_ on localization of statistical power. We repeat the data-driven simulation design, setting the maximum radius *h*_*m*_ to 5, 10, 15, 20, 25, and 30mm. Localization of power does not drastically change as we increase *h*_*m*_ from 20 to 25 or 30, and given that a smaller maximum radius is more computationally efficient, we consider our selection of a maximum radius of 20mm to be adequate.

#### 3.2.3. Comparison to spatial null modeling approaches

Despite inherent differences between permutation null and spatial null modeling methods, a comparison is warranted as the spatial null approaches have been widely adopted in studies of intermodal correspondence over the last few years [8]. Therefore, we apply the spin test and BrainSMASH to group-averaged data from each subsample of *N* =50 and 100 PNC participants considered in our simulations above for comparison. For the spatial null modeling methods, we use only 200 “permutations” (i.e., spherical rotations in the spin test and permutation of vertices in BrainSMASH in this context) due to the computational burden of running many simulations and permutations for each method. To make the spatial model refitting involved in each iteration of BrainSMASH computationally feasible to repeat across many simulations, we use parcellated brain maps with *V* = 501 regions per hemisphere, described by Schaefer et al. [29], instead of the Freesurfer vertex-level images with nearly 10,000 vertices per hemisphere.

In tests of correspondence between group-level maps of CBF and ALFF, we find that the power of the spin test is 97.7% and 100% for data averaged over *N* = 50 and *N* = 100 participants, respectively. For BrainSMASH, the power is 100% for group-averaged data in samples of both *N* = 50 and *N* = 100. However, for cortical thickness and *n*-back, there was no case that resulted in a significant *p*-value from either method in either sample size. The seemingly extreme result can be partially explained by the spin test and BrainSMASH using group-averaged maps as input, which are very similar across different samples (i.e., highly invariant to resampling different participants), because the sample means for each modality (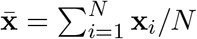 and 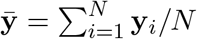) converge to the corresponding population means as *N* gets larger.

Therefore, the permutation null considered in our type I error simulations in Section 3.2 cannot be used to evaluate the type I error levels of these methods. Instead, we evaluate these methods’ type I error rates given the variability across different draws from each method’s null distribution, which are used as test statistics for computing a *p*-value under each method’s null. For the spin test, a draw from the “rotation null” hypothesis is used, so that the rotated mean maps from one modality are compared to the observed mean maps of another. We find the spin test’s type I error rates are 3.7% (95% binomial confidence interval (CI) 2.7-5.1%) for *N* = 50 and 3.6% (CI 2.6-4.9%) for *N* = 100 between CBF and ALFF, and 5.1% (CI 3.9-6.6%) for *N* = 50 and 5% (CI 3.8-6.5%) for *N* = 100 between cortical thickness and *n*-back. For BrainSMASH, a spatial autocorrelation-preserving surrogate brain map is used, and we find type I error rates of 7.9% (CI 6.4-9.7%) for *N* = 50 and 7.2% (CI 5.8-9.0%) for *N* = 100 between CBF and ALFF, and 9.1% (CI 7.5-11.0%) for *N* = 50 and 8.4% (CI 6.8-10.3%) for *N* = 100 between cortical thickness and *n*-back. Our finding of type I error rate inflation in BrainSMASH may be suggestive that, in our current implementation of BrainSMASH, the generated surrogate maps may not properly mimic the spatial structure of the observed data. Furthermore, given the variability in the spatial autocorrelation structures of different modalities (Figure 2), it is conceivable that BrainSMASH may be sensitive to the choice of which modality is used for generating surrogate maps. To our knowledge, the impact of the choice of which modality from which to generate surrogate brain maps on statistical inference and false positive rates has not yet been studied. In contrast, a strength of permutation null methods, including CLEAN-R and SPICE, is that they are insensitive to the choice of modality involved in permutations.

### 3.3. Data analysis: global testing and localization of sex differences

Finally, we apply CLEAN-R and the two other methods for testing and localizing intermodal correspondence to examine associations between CBF and ALFF—first within stratified subgroups of males (*N* = 353) and females (*N* = 478) from the PNC, and then to test and localize between-group differences (while controlling for age) using the methodology described in Section 2.3. Here, we focus on CBF/ALFF only, as cortical thickness/*n*-back did not show high levels of localized power from any method in our analysis above.

Results from sex-stratified global tests of correspondence using each method as well as maps of test statistics for localization of intermodal correspondence among males and females are shown in Figure 7(a). Un-adjusted global *p*-values for each method (shown above each set of maps) suggest that all three methods may detect significant overall correspondence between CBF and ALFF in both males and females. However, CLEAN-R provides the most interpretable maps for localizing CBF/ALFF associations, amplifying and detecting signal of intermodal correspondence even in very small regions that are undetected by the mass-univariate and spatially un-adjusted implementation of CLEAN-R.

**Figure 7:**
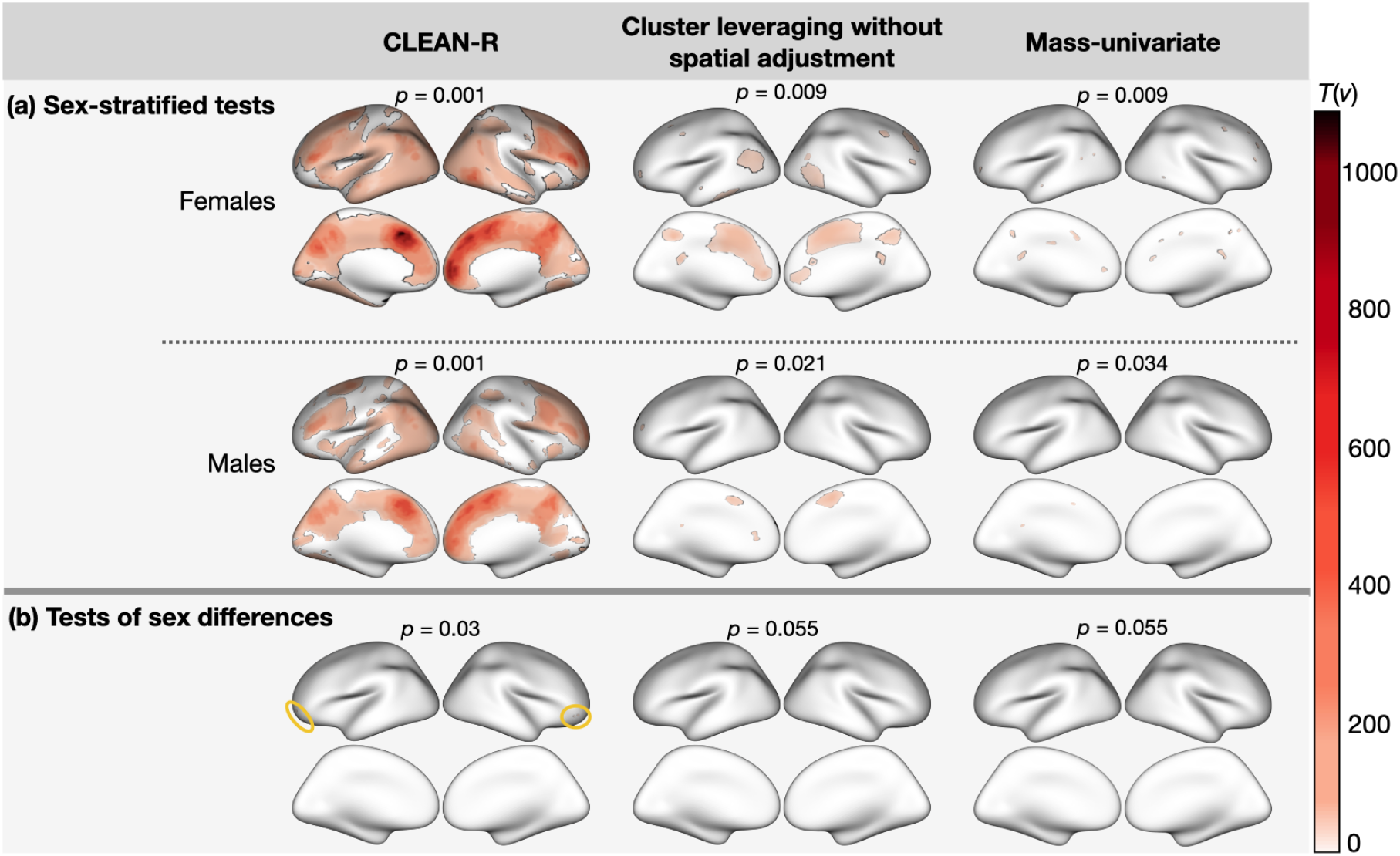
Testing and localizing intermodal correspondence between cerebral blood flow (CBF) and amplitude of low frequency fluctuations (ALFF) among *N* = 478 females and *N* = 353 males in the Philadelphia Neurodevelopmental Cohort. The *p*-values from global significance tests are provided above each map. Panel (a) includes test statistic maps from CLEAN-R, clusterwise inference without spatial adjustment, and mass-univariate tests of correspondence between CBF and ALFF applied separately in males and females. We used permutation to methods control FWER at the rate of *α* using permutation, and thresholded vertices if their statistics is less than the FWER-controlling threshold. Panel (b) includes thresholded test statistic maps from testing sex differences in correspondence between CBF and ALFF, using an extension of the original CLEAN-R methodology discussed in Section 2.3. CLEAN-R localizes significant sex differences in intermodal correspondence to four vertices (two in each hemisphere), which are encircled in yellow. Neither the clusterwise method without spatial adjustment nor the mass-univariate approach localize intermodal correspondence to any specific vertices.

In Figure 7(b), we compare sex differences in correspondence between CBF and ALFF. In global tests for overall sex differences in correspondence throughout the entire brain, CLEAN-R is the only to suggest localized correspondence between CBF and ALFF at any vertices (*p*=0.03). In a study of the same population, Baller et al. [24] evaluated sex differences in intermodal coupling [3], and using an adaptation of Alexander-Bloch et al. [6]’s spin test for testing network enrichment, found significant sex differences in CBF-ALFF coupling that were especially pronounced within Yeo et al. [10]’s frontoparietal control network. We note that two of the four significant vertices identified by CLEAN-R lie in the frontoparietal control network; however, given the small number of significant vertices, a follow-up study including a power analysis would be necessary to confirm this.

## 4. Discussion

### 4.1. Summary

In this paper, we proposed CLEAN-R, a new method for statistical inference on and localization of correspondence between two neuroimaging modalities. CLEAN-R offers insights and interpretability beyond what has been offered by previous methods for testing intermodal correspondence (e.g., global tests) by providing spatial localization of intermodal correspondence while preserving high replicability, including in relatively small samples (e.g., *N* = 50). While the idea of localization itself is a classical topic in neuroimaging (e.g., mass-univariate analyses, such as in Figure 1, and clusterwise inference for the general linear model [11, 13]), our proposed method offers greater replicability for localizing intermodal associations by accounting for modality-specific spatial autocorrelation. In practice, our method will broaden the scope of questions that can be asked and strengthen evidence derived from multimodal neuroimaging research by enhancing the statistical rigor of methods used for testing and localizing intermodal correspondence as well as testing and localizing group differences in correspondence. With a better understanding which regions of the brain are driving significant between two modalities, the interpretability and actionability of neuroimaging research will dramatically improve.

Using data from the Philadelphia Neurodevelopmental Cohort (PNC), we illustrated our method’s performance in practice in comparison to existing methods. Specifically, we showed that CLEAN-R exhibits high statistical power for detecting global correspondence between cerebral blood flow (CBF) and amplitude of low frequency fluctuations (ALFF). While the finding that CBF and ALFF are related is not new, our methodology for approaching this question offers new insight, by providing a more interpretable way to map this relationship, which might inform our understanding of which regions of the brain are driving statistically significant global tests. This aspect of intermodal correspondence testing was not part of spatial null modeling approaches [6, 7] or our previously proposed permutation method [9], since these methods involved quantifying global associations across the brain (or across pre-defined subregions), which cannot be mapped spatially.

Although we did not identify vertices with high replicability in the correspondence between cortical thickness and *n*-back, which is expected due to the sparsity of signal (Figure 1) CLEAN-R’s power was notably higher than other methods in small-sample data-driven simulation studies (Figure 5). In our earlier work [9], we also found some evidence for significant associations between *n*-back activation and cortical thickness for certain age groups when testing correspondence within functional brain networks. While other global tests require pre-defining local brain regions to detect highly localized associations, CLEAN-R’s spatially-enhanced global test statistic amplifies signal of localized associations, while controlling for false positives.

Outside the realm of intermodal correspondence analysis and testing, this paper contributes to important ongoing discussions on clusterwise inference in neuroimaging research. Expanding on ideas from Park and Fiecas [13], in our current work we argue that modeling spatial autocorrelation can mitigate the problems brought to light, for example, by Eklund et al. [14] and Noble et al. [30]. Also, compared to existing approaches that mostly focus on the spatial autocorrelation of the test statistics map [7], which could make the parametrization of spatial covariance structure intractable, our proposed approach offers a straightforward and interpretable way to account for parametric spatial Gaussian process via empirical variogram analysis.

### 4.2. Limitations and future directions

The assumption of brain-wise spatial stationarity in CLEAN-R may be a limitation, as this assumption has been subject to criticism in previous work on both clusterwise inference [14] and intermodal correspondence testing [9]. Despite the support for the assumption of stationary processes via variogram analysis within each cortical hemisphere in the context of our current study (Figure 2), such an analysis may not capture more heterogeneous covariance structures for specific subregions of the brain, as we discuss in [9]. But while the stationarity assumption warrants careful consideration, it is conceivable that even with imperfect assumptions about the spatial structure of the data, using spatially adjusted data makes aggregating information within clusters more tenable than using non-adjusted spatial data. We believe a more flexible nonstationary spatial Gaussian process would result in even higher replicability by meeting the data generating model closely.

Nonetheless, as we emphasize in Section 2.2.1, the current implementation of CLEAN-R calls for careful exploratory analyses (as in Figure 2) to determine both whether the stationarity assumption is met (e.g., based on the shape of empirical variograms) and which parametric assumption is most suitable (e.g., by comparing empirical variograms to theoretical variograms). In theory, future researchers could easily apply CLEAN-R to any set of neuroimaging modalities, including volumetric data in 3D space. But in practice, the performance of our method could be contingent on what assumptions are made, which may be context-specific. The impact of CLEAN-R’s stationarity assumption on inference remains an important question, which we look forward to investigating in the future.

Our method has room for improvement and extension. First, one possible way to circumvent the parametric assumptions would be to consider Ye et al. [31]’s non-parametric variogram model, which could more flexibly account for different sources of autocorrelation in images due not only to spatial proximity but also functional networks. Second, because the proposed clusterwise inference is limited to using spatial information only, it would also be interesting to consider alternative approaches to defining clusters, such as Noble et al. [32]’s functional network-based clusters, which improved statistical power in fMRI-based association studies. Third, it would be interesting to extend the current methodology to localize time-varying correspondence between two imaging modalities—for example, using the Adolescent Brain Cognitive Development (ABCD) study that contains a rich collection of longitudinal data from multiple modalities [33, 12]. These limitations of the current paper present numerous opportunities for future research.

### 4.3. Conclusions

CLEAN-R has numerous strengths, including its high statistical power, and in particular, its ability to spatially localize intermodal associations with high replicability in small samples, as suggested by our resampling-based simulation studies using real data. Another tremendous advantage of CLEAN-R is its computational efficiency. Spatial modeling of each modality is required only once per analysis, and a test involving thousands of permutations can be completed in a matter of a few minutes for several hundred subjects. In contrast, methods involving spatial null models (e.g. Alexander-Bloch et al. [6], Burt et al. [7]) can be much more computationally intensive by generating an empirical null distribution that involves altering high-dimensional images repeatedly. An R package and detailed documentation are publicly available on GitHub, making CLEAN-R an efficient, powerful, and user-friendly approach to test and localize intermodal correspondence.

## Acknowledgments

This research was supported by the National Science Foundation Graduate Research Fellowship Program, the National Institute of Health (R01MH123563, 2T32MH019112, R01MH107235, R01MH119219, R01NS111115, R01MH113550, R01EB022573, R01MH120482, RF1MH116920, K08MH120564, R01MH112847, R01NS060910), the National Institute of Mental Health (1ZIAMH002949), and the Natural Sciences and Engineering Research Council of Canada (RGPIN-2022-04831)

## Data and code availability statement

Raw neuroimaging data from the PNC are publicly available at the dbGaP (phs000607.v3.p2). An R package with code for conducting statistical analyses using CLEAN-R is available on GitHub at https://github.com/junjypark/CLEAN.

## CRediT authorship contribution statement

**Sarah M. Weinstein**: Conceptualization, Methodology, Software, Formal analysis, Validation, Visualization, Writing - original draft. **Simon N. Vandekar**: Supervision, Writing - review & editing. **Erica B. Baller**: Supervision, Writing - review & editing. **Danni Tu**: Methodology, Writing - review & editing. **Azeez Adebimpe**: Software, Formal analysis, Supervision, Writing - review & editing. **Tinashe M. Tapera**: Software, Formal analysis, Supervision, Writing - review & editing. **Ruben C. Gur**: Supervision, Writing - review & editing. **Raquel E. Gur**: Supervision, Writing - review & editing. **John A. Detre**: Supervision, Writing - review & editing. **Armin Raznahan**: Supervision, Writing - review & editing. **Aaron F. Alexander-Bloch**: Supervision, Writing - review & editing. **Theodore D. Satterthwaite**: Supervision, Writing - review & editing, Funding acquisition. **Russell T. Shinohara**: Methodology, Supervision, Writing - review & editing, Funding acquisition. **Jun Young Park**: Conceptualization, Methodology, Software, Supervision, Writing - review & editing, Funding acquisition.

## Suppementary Material

### S1. CLEAN-R algorithm

The following algorithm summarizes Section 2.2 in the paper.

#### Algorithm S1.1

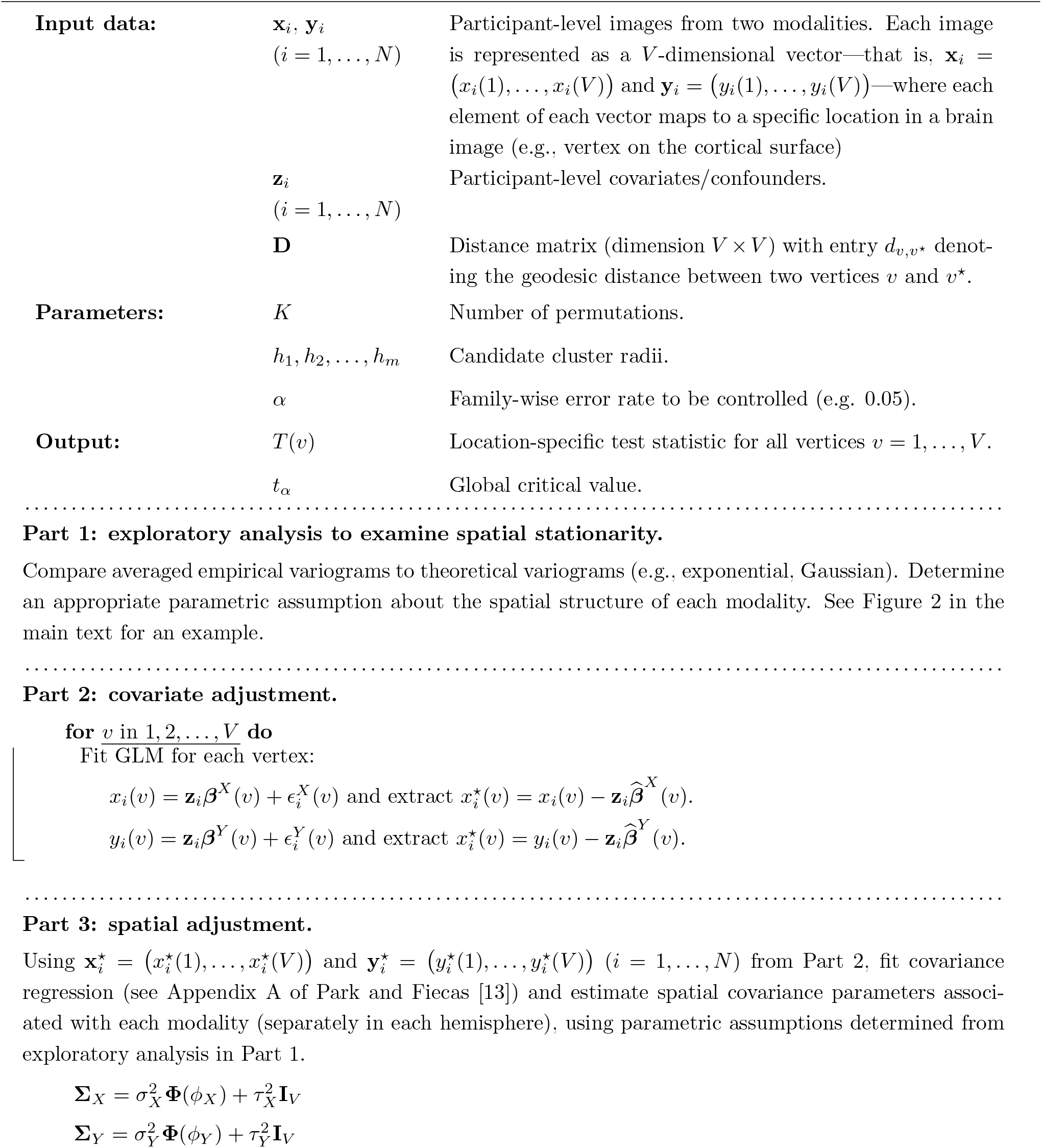
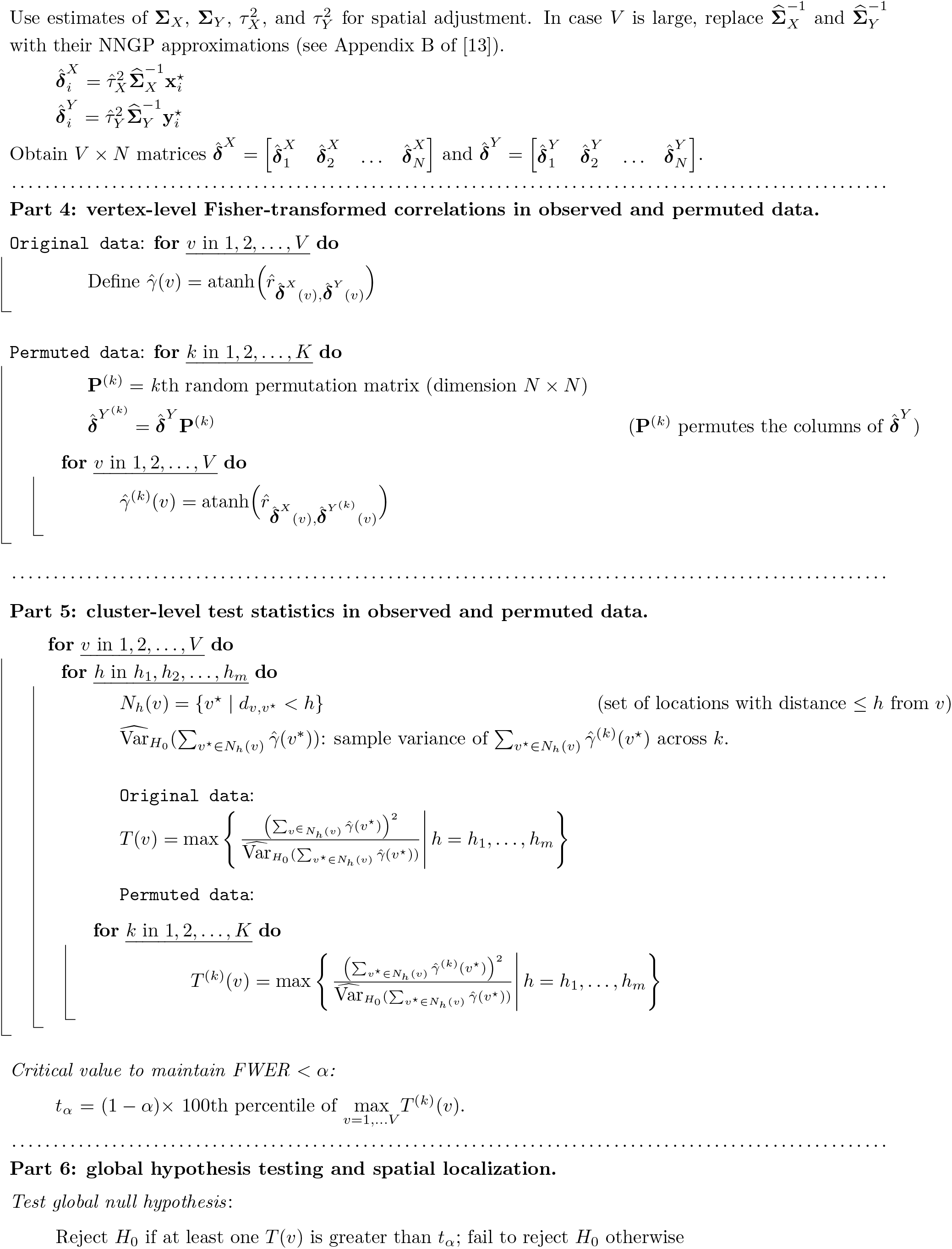
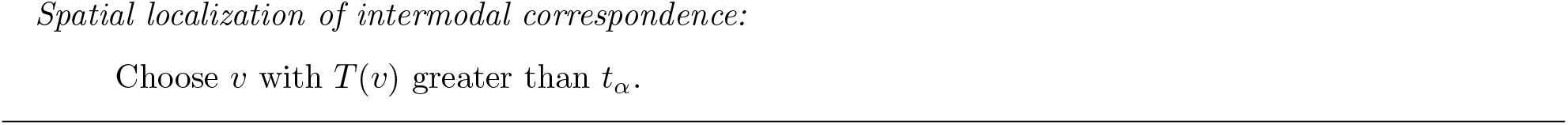

*Modifications to Algorithm 1 to implement other methods in this paper for testing & localizing intermodal correspondence*.

“Yes” and “no” in the table below express whether or not each part of the algorithm is included in implementation of these other approaches.

|  | Cluster-leveraging without spatial adjustment | Mass-univariate |
| --- | --- | --- |
| **Part 1**<br>(exploratory analysis to examine spatial stationarity) | no | no |
| **Part 2**<br>(covariate adjustment) | yes | yes |
| **Part 3**<br>(spatial adjustment) | no | no |
| **Part 4**<br>(vertex-level Fisher-transformed correlations in observed and permuted data) | yes, with modifications:<br>replace $\hat{\delta}_i^X$ with $\mathbf{x}_i^*$<br>replace $\hat{\delta}_i^Y$ with $\mathbf{y}_i^*$ | yes, with modifications:<br>replace $\hat{\delta}_i^X$ with $\mathbf{x}_i^*$<br>replace $\hat{\delta}_i^Y$ with $\mathbf{y}_i^*$ |
| **Part 5**<br>(cluster-level test statistics in observed and permuted data) | yes | yes, with modification:<br>set $h_m = 0$, so that we don't incorporate information from neighbors (i.e., cluster has radius 0). |
| **Part 6**<br>(global hypothesis testing and spatial localization) | yes | yes |

### S2. Multimodal neuroimaging data collection and processing

In this study, we analyze a subset of neuroimaging data collected as part of the Philadelphia Neurodevelopmental Cohort (PNC) [20]. PNC investigators recruited a total of *N* =9,498 children and adolescents ages 8-21. A subset of PNC participants (*N* =1,601) underwent multimodal magnetic resonance imaging (MRI) scans at the Hospital of University of Pennsylvania in the same Siemens TIM Trio 3T scanner [15], including structural MRI and resting-state and task-based functional MRI (fMRI). T1-weighted images were acquired via a magnetization-prepared, rapid acquisition gradient echo (MPRAGE) protocol, during which participants’ heads were stabilized using foam pads in an effort to reduce motion during image acquisition. For quality control purposes, image quality was rated by three experienced image analysts [34].

Cortical thickness measurements were extracted from T1-weighted structural images (with 0.9 × 0.9 ×1mm voxel resolution). Specifically, we quantified cortical thickness as the minimum distance between pial and white matter surfaces [22].

fMRI data collection included both resting-state (i.e., task-free) as well as imaging during task completion. fMRI data collection used a blood oxygen level dependent (BOLD) sequence with a single-short, interleaved multi-slice, gradient-echo, echo planar imaging sequence and a voxel resolution of 3 × 3 × 3 mm voxels over 46 slices. Preprocessing of fMRI scans included the eXtensible Connectivity Pipeline (XCP) Engine to reduce noise in images introduced by participant motion in the scanner [37].

Cerebral blood flow (CBF) and amplitude of low-frequency fluctuations (ALFF) were extracted and quantified from resting-state fMRI measurements, which were collected over the course of approximately 6 minutes. CBF measurements were quantified using parameters from arterial spin labeling (ASL) measurements (e.g., calculated as a function of the difference in signal between control and label acquisitions, the longitudinal relaxation rate of blood, and other fixed parameters). Measurements of the amplitude of low-frequency fluctuations (ALFF) were were calculated as the sum of amplitudes within the 0.01-0.08 Hertz low frequency range. Additional details on the quantification of CBF and ALFF measurements are provided by Baller et al. [24], who analyzed the same cohort of participants from the PNC.

fMRI acquisition also occurred while participants completed the *n*-back task sequence, which has been previously shown to trigger activity within the executive function network [35]. In this sequence, participants were shown various pictures sequentially on a screen. They were instructed to push a button if the stimulus they saw was identical to the *n*th previous stimulus. For example, a 2-back sequence would involve pressing the button if the current picture matched the one that appeared just before the previous one. In a “0-back” sequence, participants were instructed to simply press the button every time a new stimulus appeared (regardless of its relationship to previous stimuli). For the current study, we quantified *n*-back activation as the percentage change in fMRI measurements between the 2-back and 0-back sequences by using the FEAT tool in the FSL library [36].

Voxel-wise measurements from both structural and functional MRI data were converted into cortical surface image representations using Freesurfer (version 5.3) [21]. Measurements were projected to the fsaverage5 template, which consisted of 10,242 vertices per hemisphere for each modality (total *V* = 20, 484). However, after removing the medial wall (a byproduct of working with a surface-based representation of brain images), the number of vertices reduced to 9,354 and 9,361 in the left and right hemispheres, respectively (total *V* = 18, 715 included in analyses).

### S3. Supplementary figures

**Figure 8:**
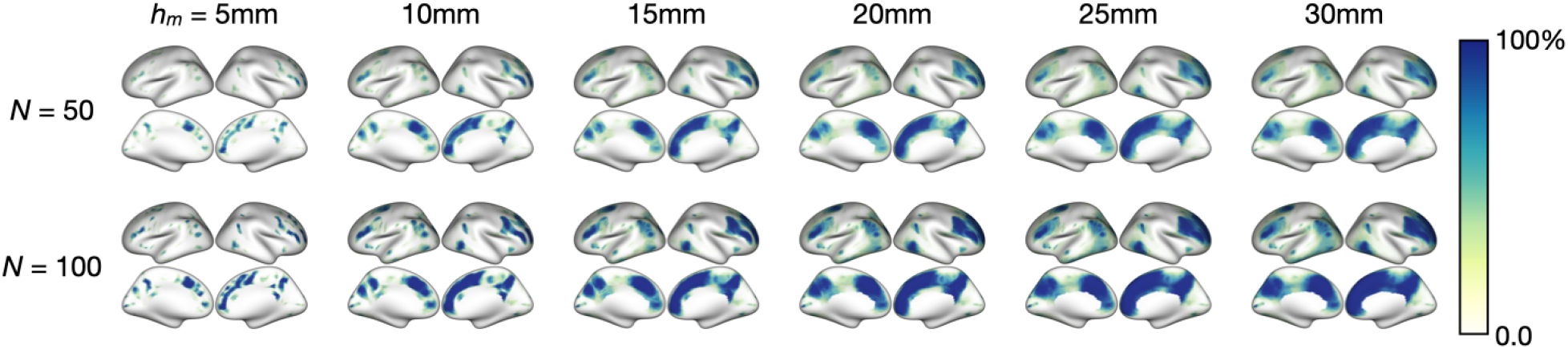
Examination of the impact of choice of maximum cluster radius (*h*_*m*_) on statistical power of CLEAN-R for localizing intermodal correspondence between cerebral blood flow (CBF) and amplitude of low frequency fluctuations (ALFF) in 1000 random subsets of *N* =50 and 100 from the Philadelphia Neurodevelopmental Cohort (PNC). Following the methods described in Sections 2.2.1 and 2.2.2, for each sub-sample of *N* =50 or 100, we compare location-statistic test statistics (*T* (*v*)) measuring correspondence between CBF and ALFF at each image location (*v* = 1, …, *V*) to *t*_*α*_, the global threshold used to determine significance (while controlling the family-wise error rate (FWER) *< α*). Similar to Figure 6 in the manuscript, at each location (*v*), statistical power is the proportion of the 1000 simulations where *T* (*v*) > *t*_*α*_.

## References

[1] R. Casanova, R. Srikanth, A. Baer, P. J. Laurienti, J. H. Burdette, S. Hayasaka, L. Flowers, F. Wood, J. A. Maldjian, Biological parametric mapping: a statistical toolbox for multimodality brain image analysis, Neuroimage 34 (2007) 137–143.

[2] X. Yang, L. Beason-Held, S. M. Resnick, B. A. Landman, Biological parametric mapping with robust and non-parametric statistics, Neuroimage 57 (2011) 423–430.

[3] S. N. Vandekar, R. T. Shinohara, A. Raznahan, R. D. Hopson, D. R. Roalf, K. Ruparel, R. C. Gur, R. E. Gur, T. D. Satterthwaite, Subject-level measurement of local cortical coupling, NeuroImage 133 (2016) 88–97.

[4] A. M. Valcarcel, K. A. Linn, S. N. Vandekar, T. D. Satterthwaite, J. Muschelli, P. A. Calabresi, D. L. Pham, M. L. Martin, R. T. Shinohara, MIMoSA: an automated method for intermodal segmentation analysis of multiple sclerosis brain lesions, Journal of Neuroimaging 28 (2018) 389–398.

[5] F. Hu, S. M. Weinstein, E. B. Baller, A. M. Valcarcel, A. Adebimpe, A. Raznahan, D. R. Roalf, T. E. Robert-Fitzgerald, V. Gonzenbach, R. C. Gur, et al., Voxel-wise intermodal coupling analysis of two or more modalities using local covariance decomposition, Human Brain Mapping (2022).

[6] A. F. Alexander-Bloch, H. Shou, S. Liu, T. D. Satterthwaite, D. C. Glahn, R. T. Shinohara, S. N. Vandekar, A. Raznahan, On testing for spatial correspondence between maps of human brain structure and function, Neuroimage 178 (2018) 540–551.

[7] J. B. Burt, M. Helmer, M. Shinn, A. Anticevic, J. D. Murray, Generative modeling of brain maps with spatial autocorrelation, NeuroImage 220 (2020) 117038.

[8] R. D. Markello, B. Misic, Comparing spatial null models for brain maps, NeuroImage 236 (2021) 118052.

[9] S. M. Weinstein, S. N. Vandekar, A. Adebimpe, T. M. Tapera, T. Robert-Fitzgerald, R. C. Gur, R. E. Gur, A. Raznahan, T. D. Satterthwaite, A. F. Alexander-Bloch, et al., A simple permutation-based test of intermodal correspondence, Human brain mapping 42 (2021) 5175–5187.

[10] B. T. Yeo, F. M. Krienen, J. Sepulcre, M. R. Sabuncu, D. Lashkari, M. Hollinshead, J. L. Roffman, J. W. Smoller, L. Zollei, J. R. Polimeni, et al., The organization of the human cerebral cortex estimated by intrinsic functional connectivity, Journal of neurophysiology (2011).

[11] S. M. Smith, T. E. Nichols, Threshold-free cluster enhancement: addressing problems of smoothing, threshold dependence and localisation in cluster inference, Neuroimage 44 (2009) 83–98.

[12] J. Y. Park, M. Fiecas, A. D. N. Initiative, et al., Permutation-based inference for spatially localized signals in longitudinal MRI data, NeuroImage 239 (2021) 118312.

[13] J. Y. Park, M. Fiecas, CLEAN: Leveraging spatial autocorrelation in neuroimaging data in clusterwise inference, Neuroimage (2022).

[14] A. Eklund, T. E. Nichols, H. Knutsson, Cluster failure: Why fMRI inferences for spatial extent have inflated false-positive rates, Proceedings of the national academy of sciences 113 (2016) 7900–7905.

[15] T. D. Satterthwaite, M. A. Elliott, K. Ruparel, J. Loughead, K. Prabhakaran, M. E. Calkins, R. Hopson, C. Jackson, J. Keefe, M. Riley, et al., Neuroimaging of the Philadelphia neurodevelopmental cohort, Neuroimage 86 (2014) 544–553.

[16] S. Banerjee, B. P. Carlin, A. E. Gelfand, Hierarchical modeling and analysis for spatial data, Chapman and Hall/CRC, 2003.

[17] C. E. McCulloch, S. R. Searle, Generalized, linear, and mixed models, John Wiley & Sons, 2004.

[18] A. Datta, S. Banerjee, A. O. Finley, A. E. Gelfand, Hierarchical nearest-neighbor gaussian process models for large geostatistical datasets, Journal of the American Statistical Association 111 (2016) 800–812.

[19] A. O. Finley, A. Datta, B. D. Cook, D. C. Morton, H. E. Andersen, S. Banerjee, Efficient algorithms for Bayesian nearest neighbor Gaussian processes, Journal of Computational and Graphical Statistics 28 (2019) 401–414.

[20] T. D. Satterthwaite, J. J. Connolly, K. Ruparel, M. E. Calkins, C. Jackson, M. A. Elliott, D. R. Roalf, R. Hopson, K. Prabhakaran, M. Behr, et al., The Philadelphia Neurodevelopmental Cohort: A publicly available resource for the study of normal and abnormal brain development in youth, Neuroimage 124 (2016) 1115–1119.

[21] B. Fischl, Freesurfer, Neuroimage 62 (2012) 774–781.

[22] A. M. Dale, B. Fischl, M. I. Sereno, Cortical surface-based analysis: I. Segmentation and surface reconstruction, Neuroimage 9 (1999) 179–194.

[23] T. Schaefer, C. Ecker, fsbrain: an R package for the visualization of structural neuroimaging data, 2020. URL: https://www.biorxiv.org/content/10.1101/2020.09.18.302935v1. xdoi:10.1101/2020.09.18.302935.

[24] E. B. Baller, A. M. Valcarcel, A. Adebimpe, A. Alexander-Bloch, Z. Cui, R. C. Gur, R. E. Gur, B. Larsen, K. A. Linn, C. M. O’Donnell, et al., Developmental coupling of cerebral blood flow and fmri fluctuations in youth (2021).

[25] S. Geuter, G. Qi, R. C. Welsh, T. D. Wager, M. A. Lindquist, Effect size and power in fmri group analysis, Biorxiv (2018) 295048.

[26] C. Lou, M. Habes, N. A. Illenberger, A. Ezzati, R. B. Lipton, P. A. Shaw, A. J. Stephens-Shields, H. Akbari, J. Doshi, C. Davatzikos, et al., Leveraging machine learning predictive biomarkers to augment the statistical power of clinical trials with baseline magnetic resonance imaging, Brain communications 3 (2021) fcab264.

[27] J. M. Franklin, S. Schneeweiss, J. M. Polinski, J. A. Rassen, Plasmode simulation for the evaluation of pharmacoepidemiologic methods in complex healthcare databases, Computational statistics & data analysis 72 (2014) 219–226.

[28] S. Benidt, D. Nettleton, Simseq: a nonparametric approach to simulation of rna-sequence datasets, Bioinformatics 31 (2015) 2131–2140.

[29] A. Schaefer, R. Kong, E. M. Gordon, T. O. Laumann, X.-N. Zuo, A. J. Holmes, S. B. Eickhoff, B. T. Yeo, Local-global parcellation of the human cerebral cortex from intrinsic functional connectivity MRI, Cerebral cortex 28 (2018) 3095–3114.

[30] S. Noble, D. Scheinost, R. T. Constable, Cluster failure or power failure? evaluating sensitivity in cluster-level inference, Neuroimage 209 (2020) 116468.

[31] J. Ye, N. A. Lazar, Y. Li, Nonparametric variogram modeling with hole effect structure in analyzing the spatial characteristics of fmri data, Journal of neuroscience methods 240 (2015) 101–115.

[32] S. Noble, M. Mejia, A. Zalesky, D. Scheinost, Leveling up: improving power in fmri by moving beyond cluster-level inference, BioRxiv (2021).

[33] N. D. Volkow, G. F. Koob, R. T. Croyle, D. W. Bianchi, J. A. Gordon, W. J. Koroshetz, E. J. Perez-Stable, W. T. Riley, M. H. Bloch, K. Conway, et al., The conception of the ABCD study: From substance use to a broad NIH collaboration, Developmental cognitive neuroscience 32 (2018) 4–7.

[34] A. F. Rosen, D. R. Roalf, K. Ruparel, J. Blake, K. Seelaus, L. P. Villa, R. Ciric, P. A. Cook, C. Davatzikos, M. A. Elliott, et al., Quantitative assessment of structural image quality, Neuroimage 169 (2018) 407–418.

[35] J. D. Ragland, B. I. Turetsky, R. C. Gur, F. Gunning-Dixon, T. Turner, L. Schroeder, R. Chan, R. E. Gur, Working memory for complex figures: an fmri comparison of letter and fractal n-back tasks., Neuropsychology 16 (2002) 370.

[36] M. Jenkinson, C. F. Beckmann, T. E. Behrens, M. W. Woolrich, S. M. Smith, Fsl, Neuroimage 62 (2012) 782–790.

[37] R. Ciric, A. F. Rosen, G. Erus, M. Cieslak, A. Adebimpe, P. A. Cook, D. S. Bassett, C. Davatzikos, D. H. Wolf, T. D. Satterthwaite, Mitigating head motion artifact in functional connectivity mri, Nature protocols 13 (2018) 2801–2826.

